# Tracing the evolution of microbial alkaline phosphatases and their role in phosphorus recycling through time

**DOI:** 10.64898/2026.06.08.730957

**Authors:** Alia Sanger, Joanne S. Boden, Andrew D. Steen, Elliott P. Mueller, Kurt O. Konhauser, Eva E. Stüeken, Rika E. Anderson, Nagissa Mahmoudi

## Abstract

Phosphorus (P) recycling in seawater is critical for maintaining nutrient availability and marine primary productivity. This process is catalyzed by alkaline phosphatases enzymes, which hydrolyze dissolved organic phosphorus (DOP) compounds, releasing inorganic P for cellular assimilation. Here, we reconstructed the evolutionary history of three major alkaline phosphatase families through deep time using phylogenetic reconciliation across the tree of life. We further quantified their distribution and cellular localization across major metabolic groups using extant genomes to assess how the ocean’s capacity for P regeneration has changed through time. Our results demonstrate that alkaline phosphatases emerged early in the Archaean, indicating that DOP has sustained marine ecosystems for most of Earth’s history. A pronounced expansion and diversification of alkaline phosphatases occurred during the Neoproterozoic, coinciding with the rise and ecological diversification of algae. Across metabolic groups, extracellular alkaline phosphatases are particularly concentrated in ferric iron reducers, fermenters, and aerobic heterotrophs, but are comparatively rare in other metabolisms. This distribution suggests that the efficiency of marine P recycling has been strongly influenced by prevailing metabolic strategies and environmental redox conditions. Overall, our results provide new insights into the enzymatic drivers of marine P cycling and the mechanisms that maintained marine productivity as Earth’s surface became progressively oxygenated and biologically complex.

## Introduction

Phosphorus (P) is an essential nutrient for all life, playing fundamental roles in the storage of genetic material (DNA and RNA), energy transfer (ATP), and cellular membranes (phospholipids). Unlike nitrogen or carbon, which can be replenished through biological fixation from atmospheric sources, the modern marine P supply is primarily delivered through riverine inputs and thus dependent on the intensity of continental weathering. Consequently, P is widely regarded as the ultimate limiting nutrient for marine primary productivity along continental margins on geological timescales (Tyrrell, 1999), and its availability is intrinsically linked to atmospheric oxygen production.

In seawater, P exists as both organic and inorganic forms that are tightly interconnected and cycle rapidly between dissolved, particulate, and biological reservoirs. The most biologically accessible form is dissolved inorganic phosphate (specifically orthophosphate, PO_4_^3-^; hereafter P_i_), which is rapidly consumed by primary producers in the photic zone and is therefore often depleted in surface waters (Michelou *et al*., 2011; Wu *et al*., 2000). However, a significant fraction of P resides in dissolved organic phosphorus (DOP) compounds, derived from phytoplankton biomass (Karl & Björkman, 2024). Marine microorganisms access much of this pool using alkaline phosphatase enzymes. These enzymes hydrolyze phosphate esters, which are estimated to comprise ∼80% of the modern seawater DOP pool (Young & Ingall, 2010), releasing P_i_ that can be readily assimilated by cells to meet cellular P demands. While alkaline phosphatases do not alter the total oceanic P inventory, they directly mediate the conversion of organic P to inorganic forms. In doing so, these enzymes regulate P regeneration, thereby exerting a strong influence on the dynamics of the marine P cycle (Karl & Björkman, 2015).

Despite their ubiquity in the modern ocean (Su *et al*., 2023), the evolutionary history of alkaline phosphatases remains poorly understood. Their diversification would have been particularly critical when global marine P concentrations were lower than today (Bjerrum & Canfield, 2002; Jones *et al*., 2015; Kipp & Stüeken, 2017; Reinhard *et al*., 2017; Hao *et al*., 2020; Planavsky *et al*., 2023; Rego *et al*., 2023). In prokaryotes, three major alkaline phosphatase enzyme families—PhoA, PhoD, and PhoX— differ in their co-factor requirements and substrate specificity (Luo *et al*., 2009), and may function intracellularly or extracellularly, depending on substrate size (Benz & Bauer, 1988) (**Fig. S1**). Here, we reconstruct the emergence and spread of alkaline phosphatases and their key regulatory genes and demonstrate that the oceans’ capacity for P recycling has shifted through time. To do this, we reconciled a time-calibrated tree of life with phylogenetic trees for alkaline phosphatase genes (*phoA, phoD, phoX)* to estimate when these genes were gained through speciation, duplication, and horizontal gene transfer. We extended this approach to a regulatory gene (*phoR*) that modulates alkaline phosphatase expression in some microorganisms (Tommassen *et al*., 1982). Finally, we quantified the distribution and cellular localization of alkaline phosphatases across major microbial metabolisms using extant genomes to infer how changing redox conditions may have shaped their prevalence in the ancient oceans.

## Results & Discussion

### An early role for DOP in sustaining Earth’s biosphere

We conducted a comprehensive survey of 865 genomes, carefully chosen to span the full phylogenetic diversity of the tree of life (Boden *et al*., 2024), in order to identify the presence of *pho*A, *pho*D, and *pho*X genes. Genes encoding alkaline phosphatases were found in 38% of the genomes we examined. The majority were found in bacterial genomes (62 of 126 bacterial phyla), but they were also present in 3 of 17 archaeal phyla and 8 of 9 eukaryotic phyla (**Figs. 1, S2**). This distribution is consistent with prior metagenomic surveys of seawater samples, which demonstrated that alkaline phosphatases are widespread among marine bacteria (Luo *et al*., 2009). Interestingly, each of the three genes were similarly distributed across genomes: *phoA* was found in 18% of the total genomes, while *phoD* and *phoX* were each present in 14% of genomes. By contrast, the regulator of the *pho* operon (*phoR*), which mediates the expression of alkaline phosphatase genes in response to low P concentrations in the environment, showed a narrow phylogenetic distribution. phoR was confined to the bacterial phylum Pseudomonadota (formerly Proteobacteria) and was present in only 4% of genomes, indicating that most microorganisms either express alkaline phosphatases constitutively or rely on alternate regulatory mechanisms to respond to P limitation.

**Figure 1.**
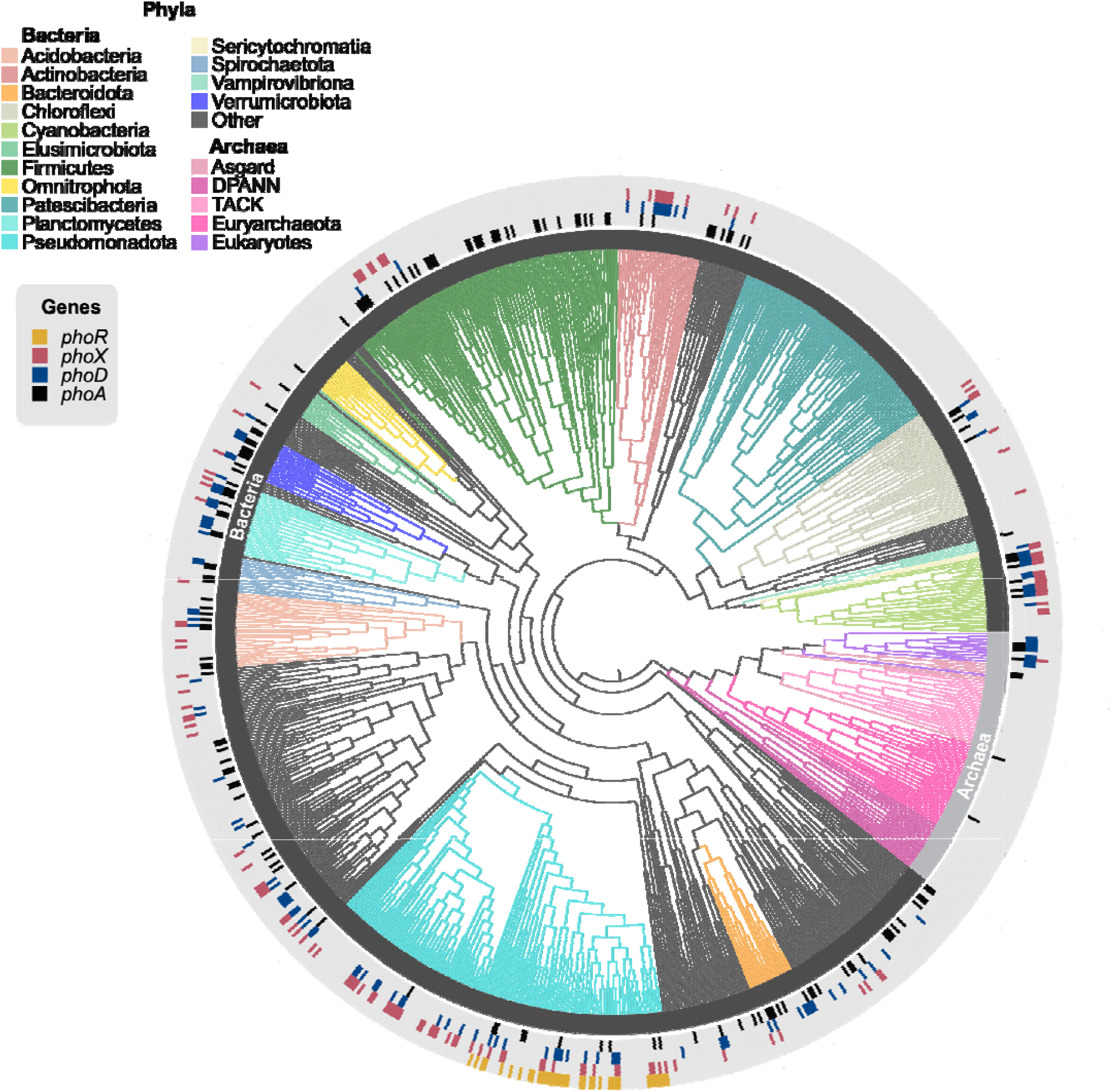
Widespread distribution of alkaline phosphatase genes in the tree of life. The branches are colored according to their phylum. The presence of the genes *phoA* (black), *phoD* (blue), *phoX* (red), and *phoR* (yellow) are represented at the tips of branches (e-value threshold 0.001). The evolutionary tree was reconstructed from 16 ribosomal proteins using a maximum likelihood approach. The associated support values are shown in Figure S2.

To reconstruct the evolutionary history of alkaline phosphatase genes, we compared protein phylogenies of the enzymes encoded by *pho*A, *phoD, pho*X, and *phoR* with our time-calibrated tree of life to infer putative events of speciation, duplication, loss, and horizontal gene transfer. We applied the autocorrelated log-normal (LN), Cox-Ingersoll-Ross (CIR), and uncorrelated gamma (UGAM) Bayesian clock models to estimate the timing for these gene events (**Fig. 2**) and provide lower bound estimates for when each gene first appeared in the phylogenetic record. We note that extinct microbial lineages leave no genomic signatures, so the actual origins of these genes could predate the earliest reported gene event. To further account for the potential variability in the relative frequency of genetic events and genome size effects, we tested a range of horizontal gene transfer event costs (6, 4, 3, 2) relative to duplication, loss, and speciation (2:1:0), with a default cost of 3 minimizing genome size variation between parent and daughter lineages over time (David & Alm, 2011) (**Figs. S3-S5**). In cases of conflicting timing estimates, we preferred results from the CIR clock model, which has been found to produce better model fit and consistency between lineages participating in the same horizontal gene transfer events (Lepage *et al*., 2007; Fournier *et al*., 2021).

**Figure 2.**
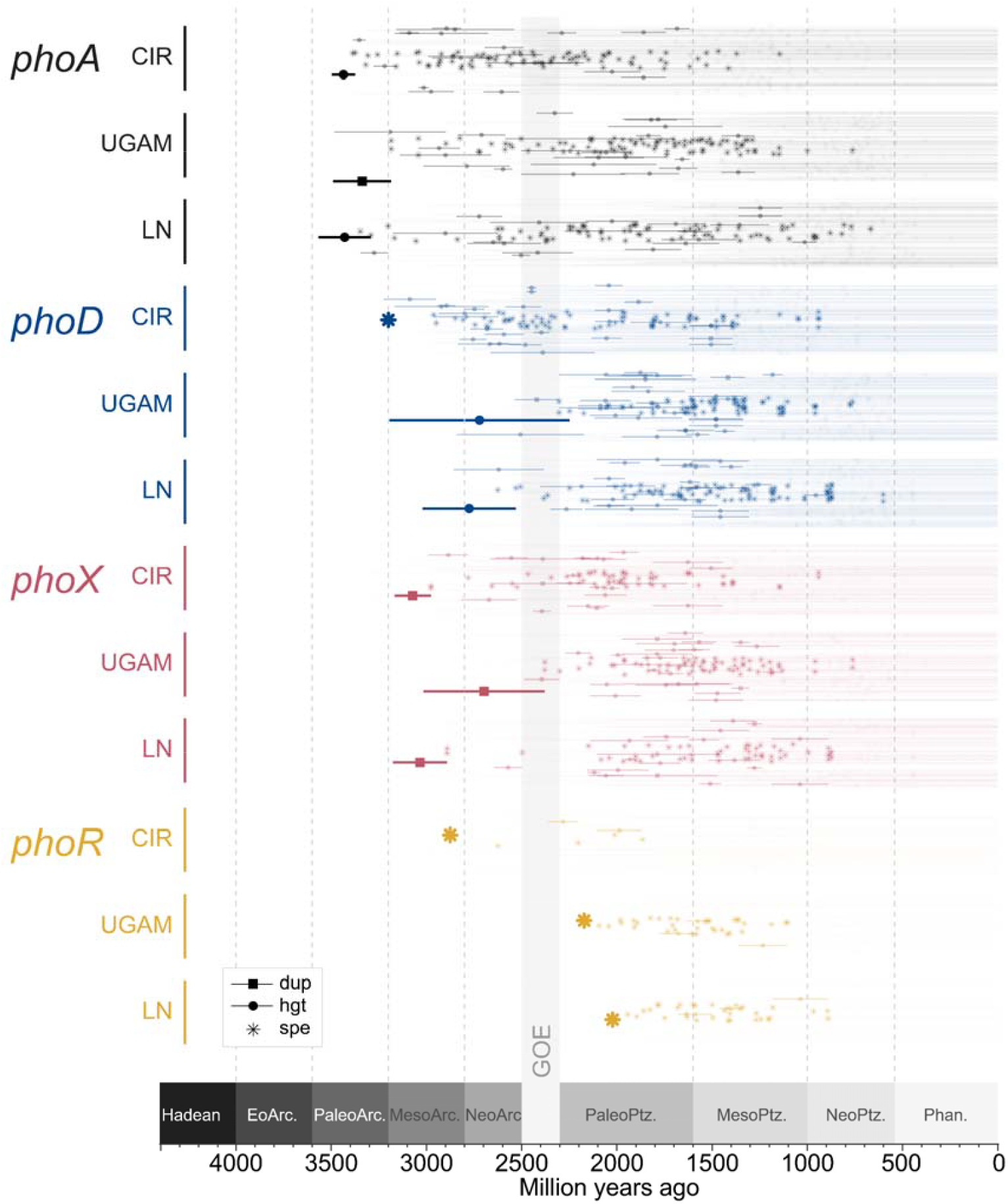
Early emergence of alkaline phosphatase genes in the Archean; *phoA* (black), *phoD* (blue) and *phoX* (red). Horizontal gene transfers (circles) and duplications (squares) of each gene are marked with colored shapes as the midpoints of lineages in which they are estimated to have occurred, depending on the results of three different clock models (CIR, UGAM and LN). Speciation events, marking a point in time where the gene was inherited from its parents, are marked with asterisks. Gene birth precedes the first inferred event (indicated by bolded symbols) for each gene. GOE refers to the Great Oxidation Event.

Our results reveal that alkaline phosphatase genes emerged very early in Earth history. *phoA* appears to be the oldest of these genes, with the earliest reconstructed event estimated to occur in the Paleoarchean, approximately 3.4 billion years ago (Ga). Both *phoD* and *phoX* follow shortly thereafter in the Mesoarchean era, spanning 3.2 Ga to 2.8 Ga (**Table S2, Fig. 2**). The early emergence of *phoA* is consistent with recent studies showing widespread distribution of *phoA* across bacterial lineages during the early stages of their evolution (Sakuma *et al*., 2025). By contrast, estimates for the emergence of the *phoR* vary between different clock models, with CIR generating earlier (Mesoarchean) estimates than UGAM and LN (Paleoproterozoic). Regardless, all analyses converge on the finding that alkaline phosphatase enzymes predate the emergence of the regulator *phoR* (**Fig. 2**). Indeed, the genomic repertoire for DOP degradation appeared to diversify early, such that multiple families of alkaline phosphatase genes were likely present across a diverse range of taxa in the Archaean (4.0-2.5 Ga) oceans. The early emergence and diversification of alkaline phosphatases imply strong selective pressures for strategies that enabled microorganisms to access DOP, thereby allowing early life to meet their P demands in marine environments where P_i_ was limited. Overall, this early proliferation of alkaline phosphatase enzymes highlights the critical, yet underappreciated, role of DOP in sustaining early microbial ecosystems and ultimately primary productivity.

### Alkaline phosphatases spread with the rise of algae

The timing and context under which alkaline phosphatases spread and diversified across the tree of life reveals ecological pressures that shaped the growth and development of Earth’s biosphere. To explore this, we quantified the occurrence of enzyme speciation events (where alkaline phosphatases are inherited vertically from the parental lineage as the parent splits into daughter lineages) as a proportion of total lineage speciation events (the number of times a parental lineage splits into daughter lineages overall at that time period, regardless of whether alkaline phosphatases are passed on in the process) through time (**Figs. 3, S6)**. By normalizing against total speciation, we can distinguish neutral patterns—where enzyme dynamics track overall diversification—from selective patterns, in which a higher proportion of lineages acquiring alkaline phosphatases would indicate evolutionary pressure favoring their use. This normalization further helps mitigate possible biases introduced by the structure of the phylogenetic tree itself, which is influenced in part by what lineages were included in constructing the tree. We focused specifically on speciation events because they can be assigned to discrete points in time, resulting in smaller uncertainties compared to gene duplications, losses, or horizontal gene transfer events, which can occur anywhere along branches that often span hundreds of millions of years. As such, we can more robustly assess the spread and diversification of alkaline phosphatases through time. This approach, in turn, provides insight into when microbial lineages inherited and retained alkaline phosphatase genes, thereby acquiring the metabolic capacity to recycle DOP.

**Figure 3.**
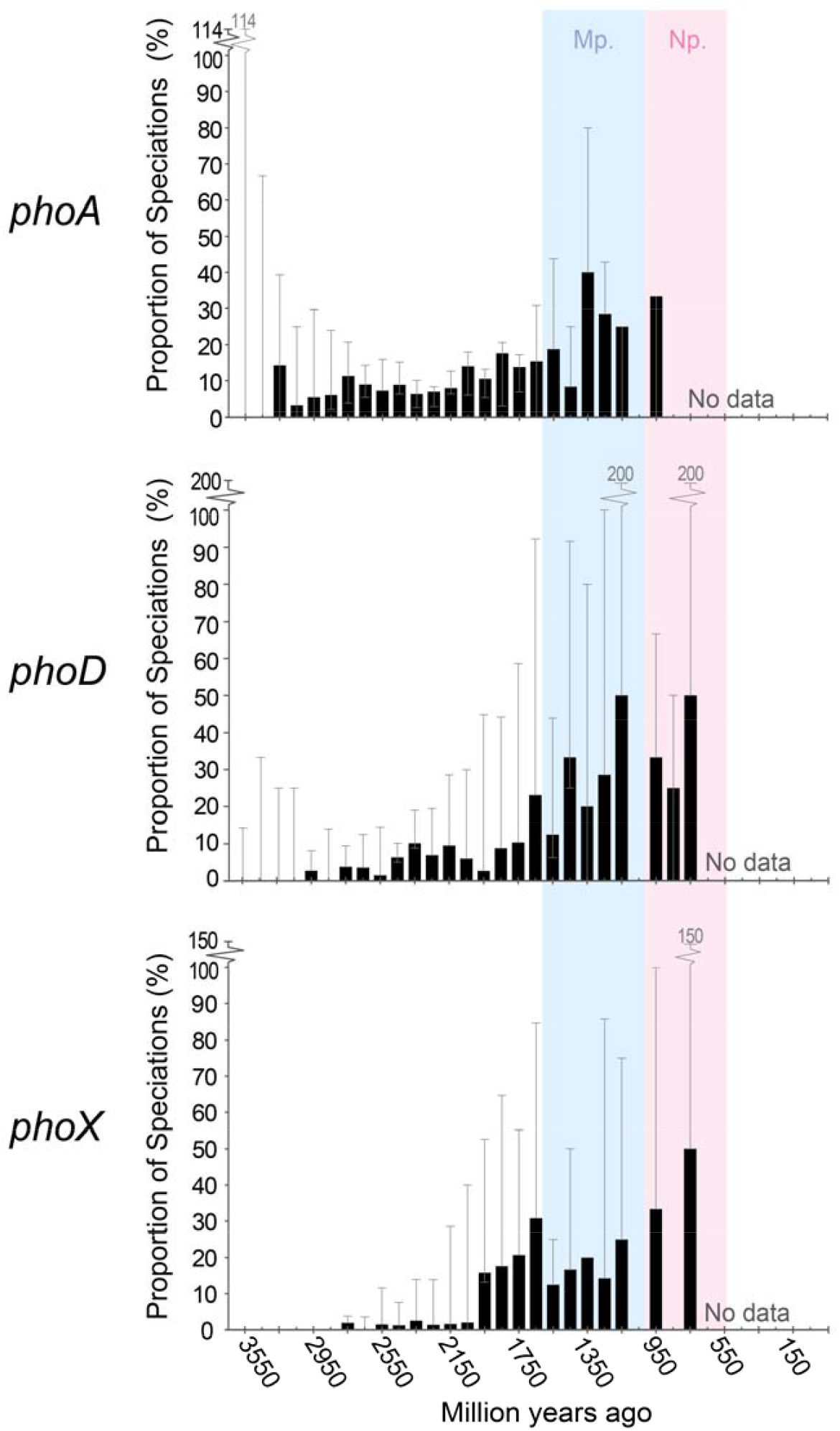
Proportional spread of alkaline phosphatase genes through time. Black bars represent the proportion of all speciation events which involved alkaline phosphatase genes calculated based on estimations made with the default HGT cost (of 3) and the Cox-Ingersoll-Ross (CIR) clock. Error bars represent the range of values estimated using HGT costs of 2, 4 and 6. Blue shading highlights the Mesoproterozoic (Mp.) and pink shading highlights the Neoproterozoic (Np.). Values on the x axis indicate the midpoint of time bins spanning 100 million years. Graphs commence at 3550 million years ago, the mid-point of the oldest time bin containing at least 3 total speciations in all clock models.

We observed a disproportionate rise in the number of speciation events involving alkaline phosphatases relative to total speciation events in the late Proterozoic, beginning in the Mesoproterozoic and peaking in the Neoproterozoic (1600 Ma and 450 Ma) (**Figs. 3, S6**). These results suggest a strong ecological pressure favouring the diversification of lineages encoding alkaline phosphatases at this time. We speculate that this expansion may have occurred in response to the broader ecological transition from Cyanobacteria-dominated to algae-dominated marine ecosystems. Photosynthetic eukaryotes are thought to have evolved through primary endosymbiosis prior to 1.6 Ga, with unambiguous fossil evidence appearing by 1.2 Ga (Sánchez-Baracaldo *et al*., 2017; Bhattacharya & Price, 2020 and references therein; Pietluch *et al*., 2025). However, it was not until the late Neoproterozoic (800-600 Ma) that algae underwent extensive diversification and rose to ecological prominence, ultimately surpassing Cyanobacteria as the dominant marine phytoplankton (Brocks *et al*., 2023).

This fundamental restructuring of marine primary producer communities likely imposed new selective pressures on P regeneration, thereby shaping the evolutionary trajectory and ecological importance of alkaline phosphatases. Compared to Cyanobacteria, algal cells are typically larger and contain larger intracellular P reservoirs bound within biomolecules such as nucleic acids and membrane lipids. Previous experimental work has shown that Cyanobacteria have relatively low lipid contents (∼11.7% on average), whereas algae are more lipid-rich (18.6–21.3%) (Finkel *et al*., 2016). This distinction between Cyanobacterial and algal cells is particularly important because a substantial fraction of the marine DOP pool is thought to be derived from membrane lipids such as phospholipid fatty acids (Clark *et al*., 1998). Consequently, the rise of algae likely expanded the size of the DOP reservoir due to a greater contribution of P-rich biomolecules. In turn, this would have increased the selective advantage of microorganisms capable of accessing DOP, thereby intensifying pressure for the proliferation and diversification of alkaline phosphatase genes at this time.

Notably, the geologic record also records a major reorganization of the marine P cycle around 800-750 Ma (Planavsky *et al*., 2023), recorded by the widespread re-appearance of phosphorites and elevated P enrichments in marine sediments (Planavsky, 2014; Reinhard *et al*., 2017). Here, our genomic record of P cycling reflects and refines these enigmatic sedimentary observations. Importantly, the expansion of alkaline phosphatase into new species does not reflect an actual increase in the total P inventory, but rather a greater capacity to recycle P from expanding and more dynamic organic pools. Because alkaline phosphatases mediate the conversion of DOP to P_i_, their expansion provides a biological mechanism for the enhanced P regeneration inferred from the geochemical record. Taken together, our findings suggest that the proliferation of alkaline phosphatases played an important role in the broader reorganization of the marine P cycle in the Neoproterozoic.

### Microbial controls on the enzymatic P regeneration

Throughout Earth’s history, the oceans have undergone profound chemical transformations in response to changing global redox conditions (Lyons *et al*., 2024). These shifts fundamentally altered nutrient availability and the distribution of electron donors and acceptors, thereby influencing which microbial metabolisms could thrive and the pathways through which key nutrients were recycled and regenerated. However, it remains unclear how the capacity to access and recycle DOP varied across different redox landscapes. To investigate this, we quantified the distribution of *phoA, phoD*, and *phoX* genes across 5,381 extant genomes categorized by metabolic groups. Unlike the tree of life used to infer the timing of gene emergence and spread, this dataset is organized by functional metabolism rather than phylogenetic representation. This framework enables a more direct assessment of how the capacity to access and recycle DOP is distributed across heterotrophic metabolisms and how that capacity may have varied under different redox regimes.

We further predicted the cellular localization of each alkaline phosphatase enzyme to estimate the fractions that are secreted into the external environment (i.e., extracellular) versus those that are retained within the cell, such as in the cytoplasm, periplasm or outer membrane (i.e., intracellular). Extracellular alkaline phosphatases are capable of hydrolyzing high-molecular-weight (MW) DOP compounds (>1000 Da) that cannot be directly taken up by cells, thereby substantially improving the ability of microorganisms to scavenge P from their external milieu by accessing a broader pool of otherwise unavailable DOP compounds (**Fig. S1**). Importantly, the ability to secrete alkaline phosphatase substantially enhances the efficiency of overall P regeneration since a large fraction of the marine DOP pool consists of large compounds exceeding 1000 Da (Karl & Björkman, 2015, 2024). This functional advantage is especially true for sedimentary environments, where much of the organic matter delivered to the seafloor occurs in the form of particles (i.e., particulate organic matter). Under these conditions in deeper waters and sediments, extracellular alkaline phosphatases play a central role in P regeneration as the primary enzymatic route by which P is liberated from organic matter.

Our results show that genes encoding intracellular alkaline phosphatases are widespread across all metabolic groups (**Fig. 4**), including both aerobes and anaerobes. This broad distribution aligns with previous work demonstrating that alkaline phosphatase enzymes are ubiquitous across diverse microbial lineages (Zimmerman *et al*., 2013). Consistent with our phylogenetic reconstructions, these findings reinforce the notion that the ability to exploit DOP to meet cellular P requirements is a deeply rooted metabolic capability that has remained essential for sustaining microbial communities for most of Earth’s history. While intracellular alkaline phosphatases were widely distributed across metabolic groups, extracellular alkaline phosphatases were more restricted, concentrated within specific metabolic groups, including dissimilatory ferric iron reducers, aerobic heterotrophs, fermenters, and dissimilatory nitrate reducers (**Fig. 4**). In contrast, sxtracellular alkaline phosphatases are nearly absent from genomes associated with dissimilatory sulfate reducers and methanogenes. This pattern closely mirrors previous observations of extracellular enzymes involved in the degradation of proteins and carbohydrates, which are also lacking from methanogens and dissimilatory sulfate reducers (Sanger *et al*., 2026). Taken together, these patterns indicate that the efficiency of DOP recycling in a given environment depends strongly on which metabolic groups are present, and by extension, the prevailing redox conditions that structure those communities.

**Figure 4.**
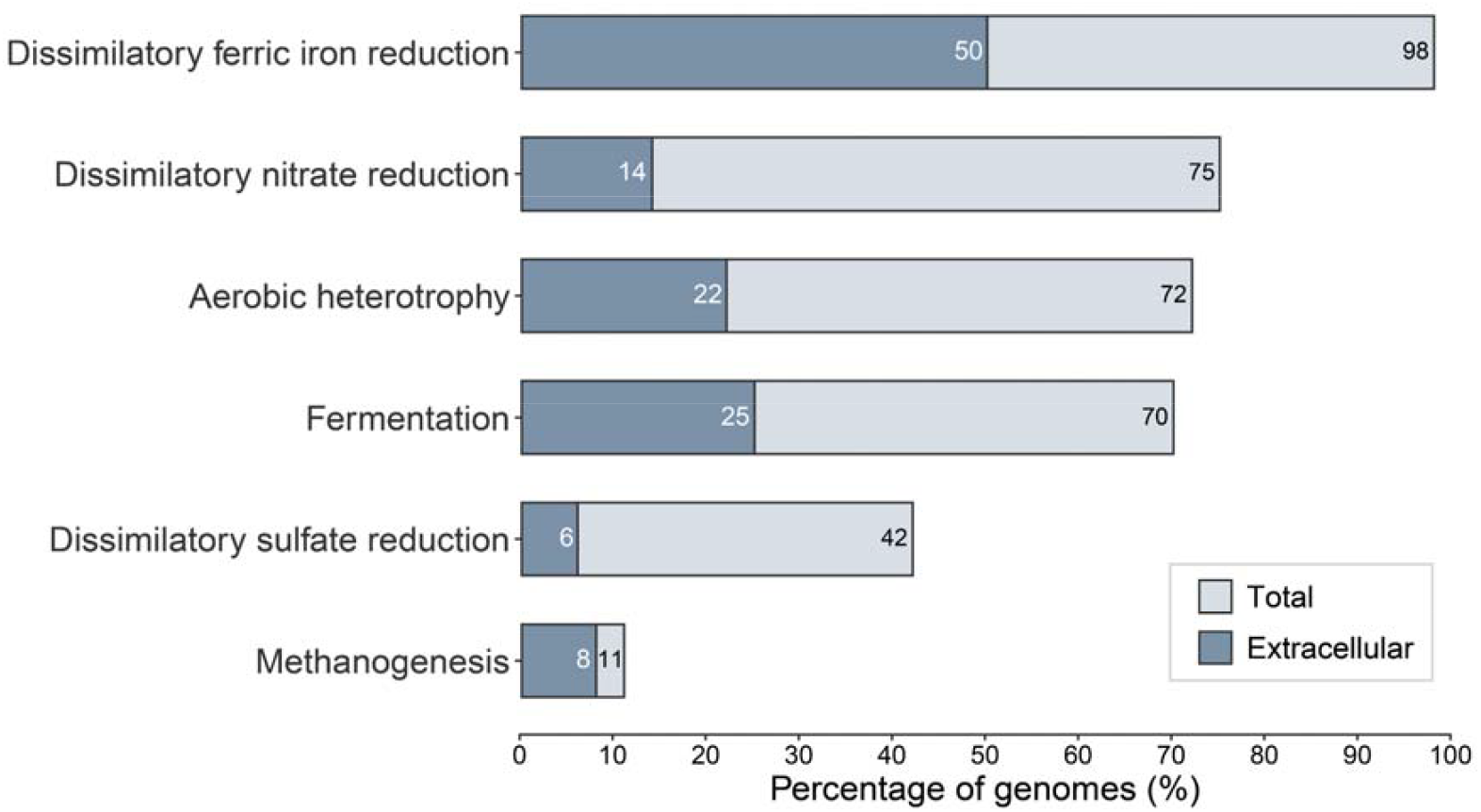
Distribution of alkaline phosphatase genes across metabolic groups. The total proportion of genomes that contain at least one alkaline phosphatase gene (*phoA, phoD*, or *phoX*; e-value threshold 0.001) is shown in light blue and the subset that contain an alkaline phosphatase gene that is predicted to localize extracellularly is shown in dark blue.

While oxygen dominates the chemistry of the modern ocean, the older Archean and early Proterozoic oceans were largely ferruginous (Konhauser *et al*., 2005; Planavsky *et al*., 2011). Following the rise of atmospheric oxygen during the Great Oxidation Event (starting around 2.45 Ga), enhanced oxidative weathering increased the delivery of P_i_ and sulfate to the oceans (Konhauser *et al*., 2011; Blättler *et al*., 2018), fueling increased primary production and widespread microbial sulfate reduction along continental margins supplied by riverine input. This led to localized euxinic conditions at continental margins, while much of the continental shelf and open ocean likely remained ferruginous, a redox structure that is thought to have persisted throughout much of the Proterozoic. Crucially, iron oxyhydroxides (e.g., ferrihydrite) efficiently adsorb organic compounds during formation, scavenging cells, their lysates, and various organic phosphorous compounds from the water column and transporting them to the seafloor (Lalonde *et al*., 2012; Li *et al*., 2021; Basinski *et al*., 2024). Aggregates of these iron oxyhydroxide-organic phosphorous compounds would have been deposited in the ferruginous deep ocean, providing an advantage for dissimilatory ferric iron reducers encoding alkaline phosphatases. Consistent with this geochemical framework, dissimilatory ferric iron reducers appear to be highly adept at degrading both low- and high-MW DOP compounds: 98% of the genomes investigated encode an alkaline phosphatase (**Fig. 5**), with 50% predicted to secrete alkaline phosphatase extracellularly. This substantial enzymatic capacity underscores the importance of DOP as a nutrient source for microorganisms inhabiting ferruginous environments, and it strongly supports the earlier notion that dissimilatory ferric iron reduction was a significant biological process in the Archean and Proterozoic iron and carbon cycles (Konhauser *et al*., 2005).

Previous geochemical models have proposed that euxinia promoted the liberation of P through the remineralization of organic matter, thereby fueling primary productivity and amplifying oxygenation of the oceans (Alcott *et al*., 2022). Our genomic data challenge this interpretation. We find that only a small fraction of dissimilatory sulfate reducers encode extracellular alkaline phosphatases that liberate P_i_ from organic matter (6% of genomes). This observation is consistent with studies showing that sulfate reducers do not directly remineralize complex organic matter but instead rely on low-molecular-weight substrates such as amino acids and acetate (Jørgensen *et al*., 2019). While euxinia would have intensified microbial sulfate reduction, our results suggest that it did not enhance enzymatic recycling of high molecular weight DOP by these microorganisms. Instead, the recycling of high molecular weight P under euxinic conditions was likely mediated by fermentative microorganisms, which exhibit a substantially higher genomic potential for extracellular alkaline phosphatase activity (25% of genomes). Yet, there is no clear mechanistic basis to expect that sulfate or sulfide concentrations would selectively stimulate DOP hydrolysis, and in turn the release of P_i_, by fermenters. Rather, under modern euxinic conditions, abiotic mineral reactions mediate P release, generating a P_i_ source for sulfate reducers. Iron minerals that strongly adsorb P_i_ (e.g., ferric oxides) react with hydrogen sulfide to form iron sulfides (Roden & Edmonds, 1997), thereby releasing previously adsorbed P_i_ into surrounding waters. This process increases local P_i_ availability independently of biological recycling. Our findings suggest that the liberation of P observed under euxinic conditions (Guilbaud, 2025) was dominated by abiotic mineral transformation, rather than by microbial recycling of DOP. This distinction reframes interpretations of Proterozoic nutrient feedbacks: although microbial sulfate reduction played a central role in shaping ocean chemistry and redox structure, the microorganisms driving P cycling were likely other heterotrophic lineages operating within these sulfidic environments.

In summary, our findings indicate that the capacity to recycle DOP, and thus regenerate P_i_, depends strongly on the dominant metabolisms present under different redox conditions. Ferruginous environments appear to favor microorganisms that have a high capacity for DOP hydrolysis, such as dissimilatory ferric iron reducers, whereas sulfate-rich euxinic settings exhibit comparatively limited biological capacity to recycle DOP. These patterns imply that major transitions in Earth’s redox landscape not only fundamentally restructured dominant metabolisms but also shifted control over P recycling within microbial communities.

## Conclusion

Alkaline phosphatases have been a fundamental component of the marine P cycle for most of Earth’s history. The early Archean origin of these enzymes suggests that DOP was an important nutrient source for some of the earliest microbial ecosystems. Their subsequent spread during the Neoproterozoic coincides with the rise of algae and evidence for intensified P recycling. Taken together, this suggests that the expansion of alkaline phosphatases at this time contributed to a fundamental reorganization of the marine P cycle. Furthermore, the uneven distribution of extracellular alkaline phosphatases across different metabolic groups demonstrates that the efficiency of P recycling depends strongly on the microorganisms favoured under different redox conditions. Collectively, the evolution and spread of alkaline phosphatases played a central role in mediating the efficiency of P regeneration, and in turn, P availability in the oceans through time.

## Acknowledgments

This work was supported by the Heising-Simons Foundation (grant no. 2024-5319 to NM). EES and JSB acknowledge funding from a NERC Frontiers grant (NE/V010824/1 awarded to EES). REA was supported as part of the Virtual Planetary Laboratory Team, which is a member of the NASA Nexus for Exoplanet System Science, and funded via the NASA Astrobiology Program ICAR Grant 80NSSC231398.

## Data Availability Statement

All data needed to evaluate the conclusions in the paper are present in the paper, Supplementary Materials and/or can be found at Figshare (https://figshare.com/s/bb3b9d5605523c8a02e3)

## Materials & Methods

### Species tree reconstruction and implementation of Bayesian molecular clocks

Our evolutionary tree of life was constructed previously (Boden *et al*., 2024) from an alignment of sixteen single-copy ribosomal proteins of 865 genomes chosen to represent the breadth of Earth’s prokaryotic microbial diversity. It is a two-domain species tree. One genome from each bacterial and archaeal phylum recognized in Parks *et al*., 2018, 2020 is included, while only 9 genomes of the 115 eukaryotic phyla described by Tedersoo, 2017 are included. This lack of Eukaryotic phyla was a conscious decision to maximise diversity of prokaryotes, which diversified around the GOE. Representative genomes were chosen based on taxonomic information from the Genome Taxonomy Database (GTDB) release 95. In this study, we implemented relaxed Bayesian molecular clocks in Phylobayes v.4.1 (Lartillot *et al*., 2009) to anchor this tree to a geological timeline. To reflect differing views on divergence time estimation, three replicates were implemented with either the lognormal (LN), cox-ingersoll-ross (CIR), or uncorrelated gamma multipliers (UGAM) clock model. A full description of the parameters is presented in Boden et al. 2024, the only difference being that here we applied an earlier calibration for the origin of methanogenesis of more than 3.46 Ga (instead of more than 2.7 Ga) based on new geochemical and biological data (Ueno *et al*., 2006; Wolfe & Fournier, 2018; Cavalazzi *et al*., 2021). Two independent chains were implemented for 129,168 to 273,674 cycles for each molecular clock model until a pair of chains reached convergence, defined as relative differences of all parameters < 0.3 and effective sizes > 50 after the first 25% of cycles had been discarded as burn-in. These relative differences and effective sizes were calculated with tracecomp implemented in Phylobayes v.4.1 (Lartillot *et al*., 2009).

### Identification of alkaline phosphatase genes

We searched in the molecular clock genomes for homologs of alkaline phosphatase genes *phoX, phoA*, and *phoD*, as well as the pho operon regulator gene *phoR*, using HMMER3 (Eddy, 2011). HMM profiles obtained from the Pfam database (*phoD*, PF09423.14; *phoX*, PF05787.17; *phoR*, PF11808.12) and NCBI’s protein family model page (*phoA*, NF007810.0) and an e-value threshold of 0.001 were applied. To ensure that the enzymes we modelled the origin of were functioning alkaline phosphatases and regulators, we took a cautious approach to identifying homologs of *phoR, phoD*, and *phoX*. This involved an extra filtration step which removed the most distantly related homologs if they did not evolve from the most recent common ancestor (MCRA) of known alkaline phosphatases or P response regulators. To do this, amino acid sequences of HMMER3 hits for each enzyme were aligned with the relevant query sequences using MAFFT v.7.4 (Katoh & Standley, 2013) with the high accuracy progressive alignment strategy (E-large-INS-1, Nakamura *et al*., 2018) designed for alignments containing multiple regions of alignable residues separated by unalignable residues and gaps (command --mpi --large --genafpair). Phylogenetically uninformative positions defined as containing more than or equal to 85 % gaps were removed using trimal v.1.4.rev22 (Capella-Gutiérrez *et al*., 2009). The resulting trimmed alignments were then used to reconstruct rooted maximum likelihood trees detailing the relationships between known enzymes and their homologs using IQTREE v.2.2.5 (Minh *et al*., 2020) with the best of four non-reversible complex substitution models (namely NQ.pfam+C20+G4, NW.pfam+C20+G4+F, NQ.bac+C20+G4,NQ.bac+C20+G4+F) chosen by ModelFinder (Kalyaanamoorthy *et al*., 2017). Branch supports were estimated by calculating ultrafast bootstraps with 1000 replicates (Hoang *et al*., 2018) and up to 5000 iterations (-bb 1000 -nm 5000). Tree topology tests, including the approximately-unbiased (AU) test (Shimodaira, 2002) were applied to measure confidence in the root placements (--root-test -zb 1000 -au). Trees were rooted on branches with the highest bootstrap support values (59.1 for PhoR, 9.26 for PhoA, 38.2 for PhoX and 77.8 for PhoD) and which passed the AU test (defined as AU p-values more than or equal to 0.1). Any homolog which did not descend from the most recent common ancestor of known enzymes was removed, resulting in a total of 35 PhoR (no homologs were removed), 183 PhoX (4 homologs were removed) and 227 PhoD (1 homolog was removed) for further analyses. Only one homolog of PhoA passed this criterion, so we kept all 200 homologs, including those which did not descend from the MCRA of the seed sequences, for further analyses. While PhoR is the most well-described alkaline phosphatase regulon, it is important to note that it is only present in a single bacterial phylum (Pseudomonadota, previously Proteobacteria) in our tree of life, which limits our interpretation of how microorganisms regulate alkaline phosphatase production in response to P_i_ concentration. There may be unidentified genes for Pho regulation present in the genomes used to construct this molecular clock.

### Evolutionary reconstruction

To estimate when *phoA, phoD, phoX*, and *phoR* evolved and spread through microbial communities, the filtered homologs were used to reconstruct phylogenetic trees for reconciliation analyses with Bayesian molecular clocks. These phylogenetic trees were reconstructed from amino acid alignments using the same strategy as above, except with a more accurate iterative alignment strategy (E-INS-i, --genafpair –-maxiterate 1000), an additional step to remove poorly sequenced residues using TAPER v.1.0.2 (Zhang *et al*., 2021), and less stringent trimming (removing alignment columns with >= 95 % gaps). These changes were made to retain as much phylogenetic signal as possible without propagating errors, following advice from Luo, 2022.

Bayesian phylogenetic trees were constructed for *phoX* and *phoR* gene trees with MrBayes v. 3.2.7a (Ronquist *et al*., 2012), employing a mixed amino acid model prior, invariant sites, and gamma-distributed site rates. Chains were considered converged when the average standard deviation of split frequencies was < 0.01, the partial scale reduction factor lay between 1.00 and 1.02, and the effective sample size scores of all parameters were > 200 after discarding the first 25% of iterations as burn in. Bayesian phylogenetic trees for *phoA* and *phoD* gene trees failed to converge within two months, so maximum-likelihood trees were constructed using IQTREE v.2.0.3 (Minh *et al*., 2020) with 1000 ultrafast bootstraps (Hoang *et al*., 2018) and one of 12 complex mixture substitution models (including LG+G4+C20+F, LG+G4+C60+F, LG+R4+C20+F, LG+R4+C60+F, Q.pfam+G4+C20+F, Q.pfam+G4+C60+F, Q.pfam+R4+C20+F, Q.pfam+R4+C60+F, Q.bac+G4+C20+F, Q.bac+G4+C60+F, Q.bac+R4+C20+F or Q.bac+R4+C60+F) with the best BIC score calculated in ModelFinder (Kalyaanamoorthy *et al*., 2017). Ultrafast bootstraps required between 200 and 1000 iterations to converge, and bootstrap trees were written to file using the -wbtl flag.

### Reconciliation with molecular clocks

Each gene tree was reconciled with the CIR clock model time-calibrated species tree using ecceTERA v. 1.2.5 (Jacox *et al*., 2016) to identify gene transfer, duplication, loss, and speciation events. To prepare the Bayesian phylogenetic trees for reconciliation we first discarded the first 25 % of the gene trees obtained from the .t files of the Mr Bayes run as burn-in. The results presented use the default event costs (horizontal gene transfer = 3, duplication = 2, loss = 1, speciation = 0, transfers to the dead allowed) and amalgamated the gene trees (amalgamate = true). The cost associated with a horizontal gene transfer event is based on the resulting impact of transfer, relative to that of duplication, loss, and speciation, on the change in genome size from the parent to the daughter lineages (David & Alm, 2011). The default ratio of 3:2:1:0 minimizes genome size variation between parents and daughter lineages over time, which is consistent with findings indicating that genome size is inherited (Martinez-Gutierrez & Aylward, 2022). However, the event cost ratio would have theoretically differed with variation in genome size, which is unconstrained for early Earth microorganisms, and there are reasons to believe that larger variation may have occurred (see discussion in Boden *et al*., 2024). Therefore, to further test how ecceTERAs, reconciliation parameters impact the estimated histories of genes related to alkaline phosphatase and determine the upper and lower bounds of our estimates, we performed reconciliations with horizontal gene transfer costs of 2,4, and 6 for all three clock models: CIR, UGAM, and LN (**Figs. S3-S5**). The symmetric median reconciliation from each gene tree was used to determine dates of gene events, confidence intervals, and divergence times using custom Python scripts, which were developed and implemented to time the evolution of sulfur and reduced P species cycling (Mateos *et al*., 2023; Boden *et al*., 2024). These scripts were applied using Python v. 2.7.5. and ensured that only reception events were counted for each occurrence of horizontal gene transfers. All trees were visualized in TreeViewer (Bianchini & Sánchez-Baracaldo, 2024).

The number of gene speciation events generally tracks the number of lineage speciation events over time (**Figs. S7-S9**). To reduce the impact of the species tree structure on the count of speciation events for alkaline phosphatases, we calculated the proportion of lineage speciation events that involve alkaline phosphatase genes relative to total speciation events. Events were grouped into 100-million-year bins, and the proportions were calculated under all three clock models (CIR, UGAM, and LN) using a horizontal gene transfer cost of 3 (**Figs. 3, S7-S9**). Horizontal gene transfer costs of 2, 4, and 6 were plotted as error bars.

### Quantification of alkaline phosphatases across metabolic groups

We used a dataset of high-quality microbial genomes across distinct metabolic groups: aerobic heterotrophy, fermentation, dissimilatory ferric iron reduction, dissimilatory sulfate reduction, dissimilatory nitrate reduction, methanogenesis. The method of dataset construction is described in Sanger *et al*., in review. Genomes were retrieved based on the presence of one or more molecular markers that corresponding to the microbial metabolisms. Molecular markers in genomes were identified using HMMER 3.3.2 (Eddy, 2011) and HMM profiles obtained from NCBI’s protein family model pages or published data from Appler *et al*., 2024 and Garber *et al*., 2020, applying the relevant bit score threshold for each HMM. For aerobic heterotrophy, we leveraged existing datasets of marine microorganisms and used a published collection (Gralka *et al*., 2023) of bacterial species isolated from coastal seawater using growth media with variable carbon substrates, and metagenome-assembled genomes (MAGs) from the GEMs catalog, Tara Oceans dataset (Tully *et al*., 2018), and Malaspina dataset (Acinas *et al*., 2021). To identify MAGs associated with aerobic heterotrophy we evaluated the presence of the complex IV (cytochrome *c* oxidase; *coxABC*), and indicator for aerobic respiration (Appler *et al*., 2024), and the absence of autotrophic and anaerobic heterotrophic molecular markers. For the *coxABC* HMM, where a sequence score cutoff was unavailable, an e-value threshold of 1×10^−50^ was used. Due to the diverse mechanisms of metabolic fermentation, which encompass a variety of start and end products, there is no molecular marker for prokaryotic fermentation. We retrieved genomes from a dataset of prokaryotic fermenters compiled by Hackmann & Zhang (2023). All genomes retained were checked for quality and completeness using CheckM v1.1.6 (Parks *et al*., 2015). Only genomes over 90% complete and with less than 5% contamination were included, meeting the high-quality level of the minimum information about a metagenome-assembled genome (MIMAG) standard (Bowers *et al*., 2017).

The resulting dataset of microbial metabolisms was queried for homologs of alkaline phosphatase genes *phoX, phoA*, and *phoD*, using HMMER3. We used HMM profiles from the Pfam database and NCBI’s protein family model page, and an e-value threshold of 1×10^−6^. The sequence hits were retrieved and were determined to code for a secreted protein based on the presence of signal peptides using SignalP 6.0 (Teufel *et al*., 2022) as described above. Before predicting subcellular localization with PSORTb, sequences were separated by predicted cell type (Gram stain) based on predicted taxonomy (Yu *et al*., 2010).

## References

Acinas SG, Sánchez P, Salazar G, Cornejo-Castillo FM, Sebastián M, Logares R, Royo-Llonch M, Paoli L, Sunagawa S, Hingamp P, Ogata H, Lima-Mendez G, Roux S, González JM, Arrieta JM, Alam IS, Kamau A, Bowler C, Raes J, Pesant S, Bork P, Agustí S, Gojobori T, Vaqué D, Sullivan MB, Pedrós-Alió C, Massana R, Duarte CM, Gasol JM (2021) Deep ocean metagenomes provide insight into the metabolic architecture of bathypelagic microbial communities. Communications Biology 4, 604.

Alcott LJ, Mills BJW, Bekker A, Poulton SW (2022) Earth’s Great Oxidation Event facilitated by the rise of sedimentary phosphorus recycling. Nature Geoscience 15, 210–215.

Appler KE, Lingford JP, Gong X, Panagiotou K, Leão P, Langwig M, Greening C, Ettema TJG, De Anda V, Baker BJ (2024) Oxygen metabolism in descendants of the archaeal-eukaryotic ancestor.

Basinski JJ, Bone SE, Klein AR, Thongsomboon W, Mitchell V, Shukle JT, Druschel GK, Thompson A, Aristilde L (2024) Unraveling iron oxides as abiotic catalysts of organic phosphorus recycling in soil and sediment matrices. Nature Communications 15, 5930.

Benz R, Bauer K (1988) Permeation of hydrophilic molecules through the outer membrane of gram-negative bacteria: Review of bacterial porins. European Journal of Biochemistry 176, 1–19.

Bhattacharya D, Price DC (2020) The Algal Tree of Life from a Genomics Perspective. In: Photosynthesis in Algae: Biochemical and Physiological Mechanisms, Advances in Photosynthesis and Respiration (eds. Larkum AWD, Grossman AR, Raven JA). Springer International Publishing, Cham, pp. 11–24.

Bianchini G, Sánchez-Baracaldo P (2024) TreeViewer: Flexible, modular software to visualise and manipulate phylogenetic trees. Ecology and Evolution 14, e10873.

Bjerrum CJ, Canfield DE (2002) Ocean productivity before about 1.9 Gyr ago limited by phosphorus adsorption onto iron oxides. Nature 417, 159–162.

Blättler CL, Claire MW, Prave AR, Kirsimäe K, Higgins JA, Medvedev PV, Romashkin AE, Rychanchik DV, Zerkle AL, Paiste K, Kreitsmann T, Millar IL, Hayles JA, Bao H, Turchyn AV, Warke MR, Lepland A (2018) Two-billion-year-old evaporites capture Earth’s great oxidation. Science 360, 320–323.

Boden JS, Zhong J, Anderson RE, Stüeken EE (2024) Timing the evolution of phosphorus-cycling enzymes through geological time using phylogenomics. Nature Communications 15, 3703.

Bowers RM, Kyrpides NC, Stepanauskas R, Harmon-Smith M, Doud D, Reddy TBK, Schulz F, Jarett J, Rivers AR, Eloe-Fadrosh EA, Tringe SG, Ivanova NN, Copeland A, Clum A, Becraft ED, Malmstrom RR, Birren B, Podar M, Bork P, Weinstock GM, Garrity GM, Dodsworth JA, Yooseph S, Sutton G, Glöckner FO, Gilbert JA, Nelson WC, Hallam SJ, Jungbluth SP, Ettema TJG, Tighe S, Konstantinidis KT, Liu W-T, Baker BJ, Rattei T, Eisen JA, Hedlund B, McMahon KD, Fierer N, Knight R, Finn R, Cochrane G, Karsch-Mizrachi I, Tyson GW, Rinke C, Lapidus A, Meyer F, Yilmaz P, Parks DH, Murat Eren A, Schriml L, Banfield JF, Hugenholtz P, Woyke T (2017) Minimum information about a single amplified genome (MISAG) and a metagenome-assembled genome (MIMAG) of bacteria and archaea. Nature Biotechnology 35, 725–731.

Brocks JJ, Nettersheim BJ, Adam P, Schaeffer P, Jarrett AJM, Güneli N, Liyanage T, Van Maldegem LM, Hallmann C, Hope JM (2023) Lost world of complex life and the late rise of the eukaryotic crown. Nature 618, 767–773.

Capella-Gutiérrez S, Silla-Martínez JM, Gabaldón T (2009) trimAl: a tool for automated alignment trimming in large-scale phylogenetic analyses. Bioinformatics 25, 1972–1973.

Cavalazzi B, Lemelle L, Simionovici A, Cady SL, Russell MJ, Bailo E, Canteri R, Enrico E, Manceau A, Maris A, Salomé M, Thomassot E, Bouden N, Tucoulou R, Hofmann A (2021) Cellular remains in a ∼3.42-billion-year-old subseafloor hydrothermal environment. Science Advances 7, eabf3963.

Clark LL, Ingall ED, Benner R (1998) Marine phosphorus is selectively remineralized. Nature 393, 426–426.

David LA, Alm EJ (2011) Rapid evolutionary innovation during an Archaean genetic expansion. Nature 469, 93–96.

Eddy SR (2011) Accelerated Profile HMM Searches. PLOS Computational Biology 7, e1002195.

Finkel ZV, Follows MJ, Liefer JD, Brown CM, Benner I, Irwin AJ (2016) Phylogenetic Diversity in the Macromolecular Composition of Microalgae. PLOS ONE 11, e0155977.

Fournier GP, Moore KR, Rangel LT, Payette JG, Momper L, Bosak T (2021) The Archean origin of oxygenic photosynthesis and extant cyanobacterial lineages. Proceedings of the Royal Society B: Biological Sciences 288, 20210675.

Garber AI, Nealson KH, Okamoto A, McAllister SM, Chan CS, Barco RA, Merino N (2020) FeGenie: A Comprehensive Tool for the Identification of Iron Genes and Iron Gene Neighborhoods in Genome and Metagenome Assemblies. Frontiers in Microbiology 11, 37.

Gralka M, Pollak S, Cordero OX (2023) Genome content predicts the carbon catabolic preferences of heterotrophic bacteria. Nature Microbiology 8, 1799–1808.

Guilbaud R (2025) Proterozoic evolution of the phosphorus cycle: Was it high or was it low? In: Treatise on Geochemistry. Elsevier, pp. 153–175.

Hackmann TJ, Zhang B (2023) The phenotype and genotype of fermentative prokaryotes. Science Advances 9, eadg8687.

Hao J, Knoll AH, Huang F, Schieber J, Hazen RM, Daniel I (2020) Cycling phosphorus on the Archean Earth: Part II. Phosphorus limitation on primary production in Archean ecosystems. Geochimica et Cosmochimica Acta 280, 360–377.

Hoang DT, Chernomor O, Von Haeseler A, Minh BQ, Vinh LS (2018) UFBoot2: Improving the Ultrafast Bootstrap Approximation. Molecular Biology and Evolution 35, 518–522.

Jacox E, Chauve C, Szöllősi GJ, Ponty Y, Scornavacca C (2016) ecceTERA: comprehensive gene tree-species tree reconciliation using parsimony. Bioinformatics 32, 2056–2058.

Jones C, Nomosatryo S, Crowe SA, Bjerrum CJ, Canfield DE (2015) Iron oxides, divalent cations, silica, and the early earth phosphorus crisis. Geology 43, 135–138.

Jørgensen BB, Findlay AJ, Pellerin A (2019) The Biogeochemical Sulfur Cycle of Marine Sediments. Frontiers in Microbiology 10, 849.

Kalyaanamoorthy S, Minh BQ, Wong TKF, Von Haeseler A, Jermiin LS (2017) ModelFinder: fast model selection for accurate phylogenetic estimates. Nature Methods 14, 587–589.

Karl DM, Björkman KM (2015) Chapter 5 - Dynamics of Dissolved Organic Phosphorus. In: Biogeochemistry of Marine Dissolved Organic Matter (Second Edition) (eds. Hansell DA, Carlson CA). Academic Press, Boston, pp. 233–334.

Karl DM, Björkman KM (2024) Chapter 9 - Biogeochemistry of organic phosphorus in the sea: Advances, challenges, and opportunities. In: Biogeochemistry of Marine Dissolved Organic Matter (Third Edition) (eds. Hansell DA, Carlson CA). Academic Press, pp. 405–482.

Katoh K, Standley DM (2013) MAFFT Multiple Sequence Alignment Software Version 7: Improvements in Performance and Usability. Molecular Biology and Evolution 30, 772–780.

Kipp MA, Stüeken EE (2017) Biomass recycling and Earth’s early phosphorus cycle. Science Advances 3, eaao4795.

Konhauser KO, Lalonde SV, Planavsky NJ, Pecoits E, Lyons TW, Mojzsis SJ, Rouxel OJ, Barley ME, Rosìere C, Fralick PW, Kump LR, Bekker A (2011) Aerobic bacterial pyrite oxidation and acid rock drainage during the Great Oxidation Event. Nature 478, 369–373.

Konhauser KO, Newman DK, Kappler A (2005) The potential significance of microbial Fe(III) reduction during deposition of Precambrian banded iron formations. Geobiology 3, 167–177.

Lalonde K, Mucci A, Ouellet A, Gélinas Y (2012) Preservation of organic matter in sediments promoted by iron. Nature 483, 198–200.

Lartillot N, Lepage T, Blanquart S (2009) PhyloBayes 3: a Bayesian software package for phylogenetic reconstruction and molecular dating. Bioinformatics 25, 2286–2288.

Lepage T, Bryant D, Philippe H, Lartillot N (2007) A General Comparison of Relaxed Molecular Clock Models. Molecular Biology and Evolution 24, 2669–2680.

Li Y, Sutherland BR, Gingras MK, Owttrim GW, Konhauser KO (2021) A novel approach to investigate the deposition of (bio)chemical sediments: The sedimentation velocity of cyanobacteria–ferrihydrite aggregates. Journal of Sedimentary Research 91, 390–398.

Luo H (ed.) (2022) Environmental Microbial Evolution: Methods and Protocols. Methods in Molecular Biology. Springer US, New York, NY.

Luo H, Benner R, Long RA, Hu J (2009) Subcellular localization of marine bacterial alkaline phosphatases. Proceedings of the National Academy of Sciences 106, 21219–21223.

Lyons TW, Tino CJ, Fournier GP, Anderson RE, Leavitt WD, Konhauser KO, Stüeken EE (2024) Co-evolution of early Earth environments and microbial life. Nature Reviews Microbiology 22, 572–586.

Martinez-Gutierrez CA, Aylward FO (2022) Genome size distributions in bacteria and archaea are strongly linked to evolutionary history at broad phylogenetic scales. PLOS Genetics 18, e1010220.

Mateos K, Chappell G, Klos A, Le B, Boden J, Stüeken E, Anderson R (2023) The evolution and spread of sulfur cycling enzymes reflect the redox state of the early Earth. Science Advances 9, eade4847.

Michelou VK, Lomas MW, Kirchman DL (2011) Phosphate and adenosine-5’-triphosphate uptake by cyanobacteria and heterotrophic bacteria in the Sargasso Sea. Limnology and Oceanography 56, 323–332.

Minh BQ, Schmidt HA, Chernomor O, Schrempf D, Woodhams MD, Von Haeseler A, Lanfear R (2020) IQ-TREE 2: New Models and Efficient Methods for Phylogenetic Inference in the Genomic Era. Molecular Biology and Evolution 37, 1530–1534.

Nakamura T, Yamada KD, Tomii K, Katoh K (2018) Parallelization of MAFFT for large-scale multiple sequence alignments. Bioinformatics 34, 2490–2492.

Parks DH, Chuvochina M, Chaumeil P-A, Rinke C, Mussig AJ, Hugenholtz P (2020) A complete domain-to-species taxonomy for Bacteria and Archaea. Nature Biotechnology 38, 1079–1086.

Parks DH, Chuvochina M, Waite DW, Rinke C, Skarshewski A, Chaumeil P-A, Hugenholtz P (2018) A standardized bacterial taxonomy based on genome phylogeny substantially revises the tree of life. Nature Biotechnology 36, 996–1004.

Parks DH, Imelfort M, Skennerton CT, Hugenholtz P, Tyson GW (2015) CheckM: assessing the quality of microbial genomes recovered from isolates, single cells, and metagenomes. Genome Research 25, 1043–1055.

Pietluch F, Mackiewicz P, Sidorczuk K, Gagat P (2025) Dating the Origin and Spread of Plastids and Chromatophores. International Journal of Molecular Sciences 26, 5569.

Planavsky NJ (2014) The elements of marine life. Nature Geoscience 7, 855–856.

Planavsky NJ, Asael D, Rooney AD, Robbins LJ, Gill BC, Dehler CM, Cole DB, Porter SM, Love GD, Konhauser KO, Reinhard CT (2023) A sedimentary record of the evolution of the global marine phosphorus cycle. Geobiology 21, 168–174.

Planavsky NJ, McGoldrick P, Scott CT, Li C, Reinhard CT, Kelly AE, Chu X, Bekker A, Love GD, Lyons TW (2011) Widespread iron-rich conditions in the mid-Proterozoic ocean. Nature 477, 448–451.

Rego ES, Busigny V, Lalonde SV, Rossignol C, Babinski M, Philippot P (2023) Low-phosphorus concentrations and important ferric hydroxide scavenging in Archean seawater. PNAS Nexus 2, pgad025.

Reinhard CT, Planavsky NJ, Gill BC, Ozaki K, Robbins LJ, Lyons TW, Fischer WW, Wang C, Cole DB, Konhauser KO (2017) Evolution of the global phosphorus cycle. Nature 541, 386–389.

Roden EE, Edmonds JW (1997) Phosphate mobilization in iron-rich anaerobic sediments: microbial Fe (III) oxide reduction versus iron-sulfide formation. Archiv für Hydrobiologie 139, 347–378.

Ronquist F, Teslenko M, Van Der Mark P, Ayres DL, Darling A, Höhna S, Larget B, Liu L, Suchard MA, Huelsenbeck JP (2012) MrBayes 3.2: Efficient Bayesian Phylogenetic Inference and Model Choice Across a Large Model Space. Systematic Biology 61, 539–542.

Sakuma M, Konno N, Gholipour S, Chen JZ, Tokuriki N (2025) Widespread promiscuous alkaline phosphatases underscore early microbial phosphite utilization.

Sánchez-Baracaldo P, Raven JA, Pisani D, Knoll AH (2017) Early photosynthetic eukaryotes inhabited low-salinity habitats. Proceedings of the National Academy of Sciences 114.

Shimodaira H (2002) An Approximately Unbiased Test of Phylogenetic Tree Selection. Systematic Biology 51, 492–508.

Su B, Song X, Duhamel S, Mahaffey C, Davis C, Ivančić I, Liu J (2023) A dataset of global ocean alkaline phosphatase activity. Scientific Data 10, 205.

Tedersoo L (2017) Proposal for practical multi-kingdom classification of eukaryotes based on monophyly and comparable divergence time criteria.

Teufel F, Almagro Armenteros JJ, Johansen AR, Gíslason MH, Pihl SI, Tsirigos KD, Winther O, Brunak S, Von Heijne G, Nielsen H (2022) SignalP 6.0 predicts all five types of signal peptides using protein language models. Nature Biotechnology 40, 1023–1025.

Tommassen J, De Geus P, Lugtenberg B, Hackett J, Reeves P (1982) Regulation of the pho regulon of Escherichia coli K-12. Journal of Molecular Biology 157, 265–274.

Tully BJ, Graham ED, Heidelberg JF (2018) The reconstruction of 2,631 draft metagenome-assembled genomes from the global oceans. Scientific Data 5, 170203.

Tyrrell T (1999) The relative influences of nitrogen and phosphorus on oceanic primary production. Nature 400, 525–531.

Ueno Y, Yamada K, Yoshida N, Maruyama S, Isozaki Y (2006) Evidence from fluid inclusions for microbial methanogenesis in the early Archaean era. Nature 440, 516–519.

Wolfe JM, Fournier GP (2018) Horizontal gene transfer constrains the timing of methanogen evolution. Nature Ecology & Evolution 2, 897–903.

Young CL, Ingall ED (2010) Marine Dissolved Organic Phosphorus Composition: Insights from Samples Recovered Using Combined Electrodialysis/Reverse Osmosis. Aquatic Geochemistry 16, 563–574.

Yu NY, Wagner JR, Laird MR, Melli G, Rey S, Lo R, Dao P, Sahinalp SC, Ester M, Foster LJ, Brinkman FSL (2010) PSORTb 3.0: improved protein subcellular localization prediction with refined localization subcategories and predictive capabilities for all prokaryotes. Bioinformatics 26, 1608–1615.

Zhang C, Zhao Y, Braun EL, Mirarab S (2021) TAPER: Pinpointing errors in multiple sequence alignments despite varying rates of evolution. Methods in Ecology and Evolution 12, 2145–2158.

Zimmerman AE, Martiny AC, Allison SD (2013) Microdiversity of extracellular enzyme genes among sequenced prokaryotic genomes. The ISME Journal 7, 1187–1199.

