## Supporting Information for "Tracing the evolution of microbial alkaline phosphatases and their role in phosphorus recycling through time"

**Table S1.** Molecular clock estimates of the emergence of crown bacteria, crown archaea, and LUCA, under three different clock models.

| Clock model | Bacteria / Ga | Archaea / Ga | LUCA / Ga |
|---|---|---|---|
| CIR | 3.43 (3.73-3.54) | 3.71 (3.81-3.62) | 4.36 (4.40-4.33) |
| UGAM | 4.02 (4.38-3.67) | 3.73 (3.83-3.62) | 4.38 (4.40-4.35) |
| LN | 3.80 (3.92-3.68) | 3.75 (3.83-3.66) | 4.38 (4.40-4.37) |

Mean (95% confidence interval). CIR = Cox-Ingersoll-Ross; LN = log-normal; UGAM = uncorrelated gamma multiplier

**Table S2.** Estimated dates for the earliest events during the evolution of alkaline phosphatase genes (*phoA*, *phoD*, *phoX*) and the P response regulator (*phoR*). Asterisks (*) indicate that the earliest event was a speciation. For these, we represent the median date of the node experiencing speciation alongside the upper and lower confidence intervals in brackets. Dates without asterisks indicate that the first event was estimated to be a horizontal gene transfer or duplication. For these, we present the midpoint of the lineage which experienced the event alongside the upper confidence interval of the start of the lineage and the lower confidence interval of the end of the lineage in brackets. Earliest events were defined as those with the oldest median age of speciation or the oldest midpoint age of lineages experiencing transfer or duplication.

| Clock Model | HGT Cost | Date of First Event / Ga | | | |
|---|---|---|---|---|---|
| | | **phoA** | **phoD** | **phoX** | **phoR** |
| CIR | 2 | 3.028 (3.158-2.904)* | 3.200 (3.324-3.092)* | 3.072 (3.271-2.855) | 2.510 (2.647-2.357)* |
| | **3** | **3.436 (3.644-3.237)** | **3.200 (3.324-3.092)*** | **3.072 (3.271-2.855)** | **2.875 (3.032-2.706)*** |
| | 4 | 3.609 (3.737-3.535)* | 3.200 (3.324-3.092)* | 3.072 (3.271-2.855) | 2.608 (2.731-2.476)* |
| | 6 | 3.995 (4.399-3.535) | 3.568 (3.689-3.498)* | 3.072 (3.271-2.855) | 2.757 (2.877-2.636)* |
| UGAM | 2 | 2.659 (2.980-2.341)* | 2.721 (3.418-1.590) | 2.697 (3.277-1.490) | 2.172 (2.339-1.985)* |
| | **3** | **3.338 (3.990-2.775)** | **2.721 (3.418-1.590)** | **2.697 (3.277-1.490)** | **2.172 (2.339-1.985)*** |
| | 4 | 4.148 (4.378-3.665)* | 2.975 (3.394-2.459) | 2.697 (3.277-1.490) | 2.172 (2.339-1.985)* |
| | 6 | 4.267 (4.400-3.665) | 4.015 (4.262-3.581)* | 3.046 (3.259-2.778)* | 2.434 (2.627-2.220)* |
| LN | 2 | 3.292 (3.543-3.018)* | 2.621 (3.043-2.100) | 3.033 (3.293-2.725) | 1.888 (2.062-1.714)* |
| | **3** | **3.429 (3.753-3.018)** | **2.776 (3.162-2.298)** | **3.033 (3.293-2.725)** | **2.023 (2.163-1.885)*** |
| | 4 | 3.803 (3.921-3.675)* | 2.953 (3.207-2.660) | 3.033 (3.293-2.725) | 2.023 (2.163-1.885)* |
| | 6 | 4.097 (4.400-3.675) | 3.723 (3.846-3.597)* | 3.294 (3.430-3.166)* | 2.253 (2.397-2.115)* |

HGT = horizontal gene transfer; CIR = Cox-Ingersoll-Ross CIR; LN = log-normal; UGAM = uncorrelated gamma multiplier

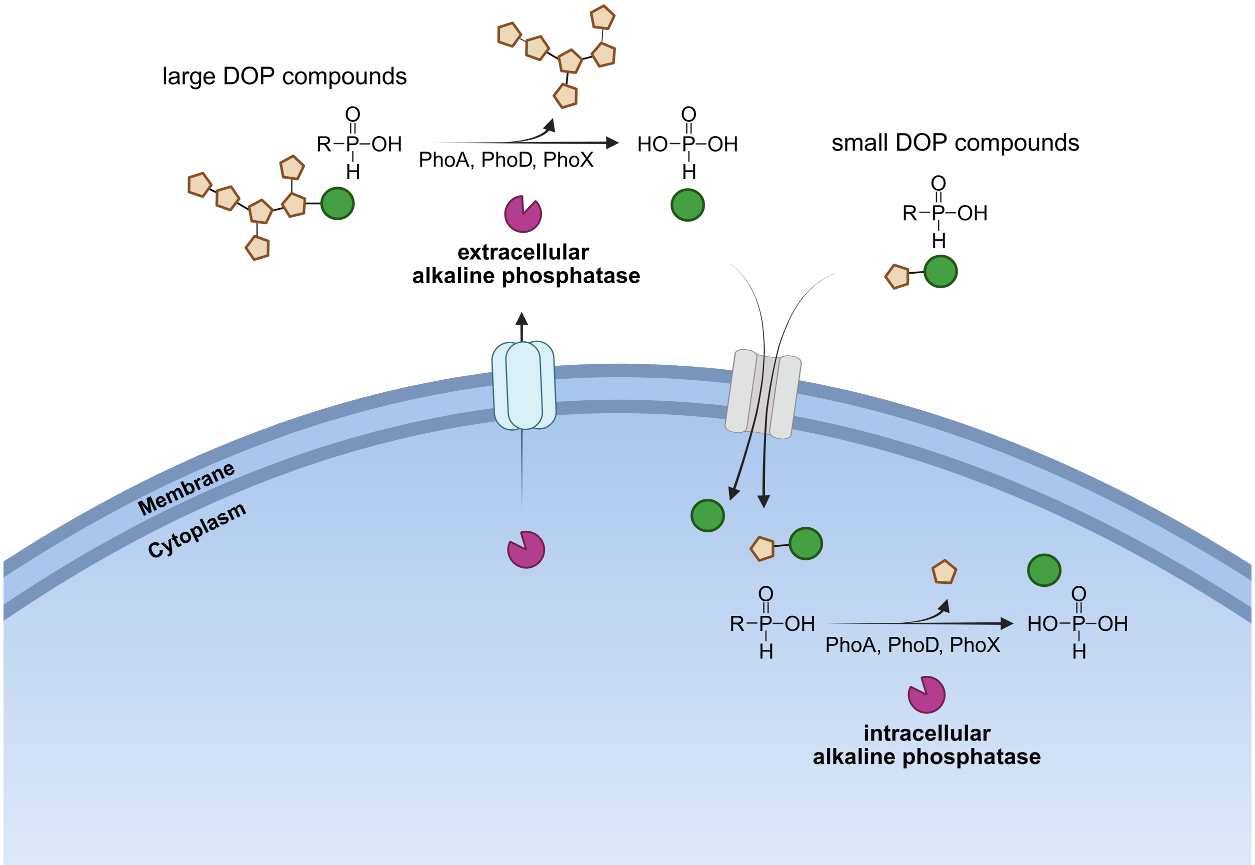


**Figure S1.** Simplified schematic depicting intracellular and extracellular alkaline phosphatases (pink). Phosphate groups are represented by green circles, and yellow hexagons represent organic matter. The cell (blue) can directly import small, low-molecular-weight (<1000 Da, Benz & Bauer, 1988) DOP compounds. To access large, high-molecular-weight DOP compounds cells must secrete alkaline phosphatases into the extracellular environment.

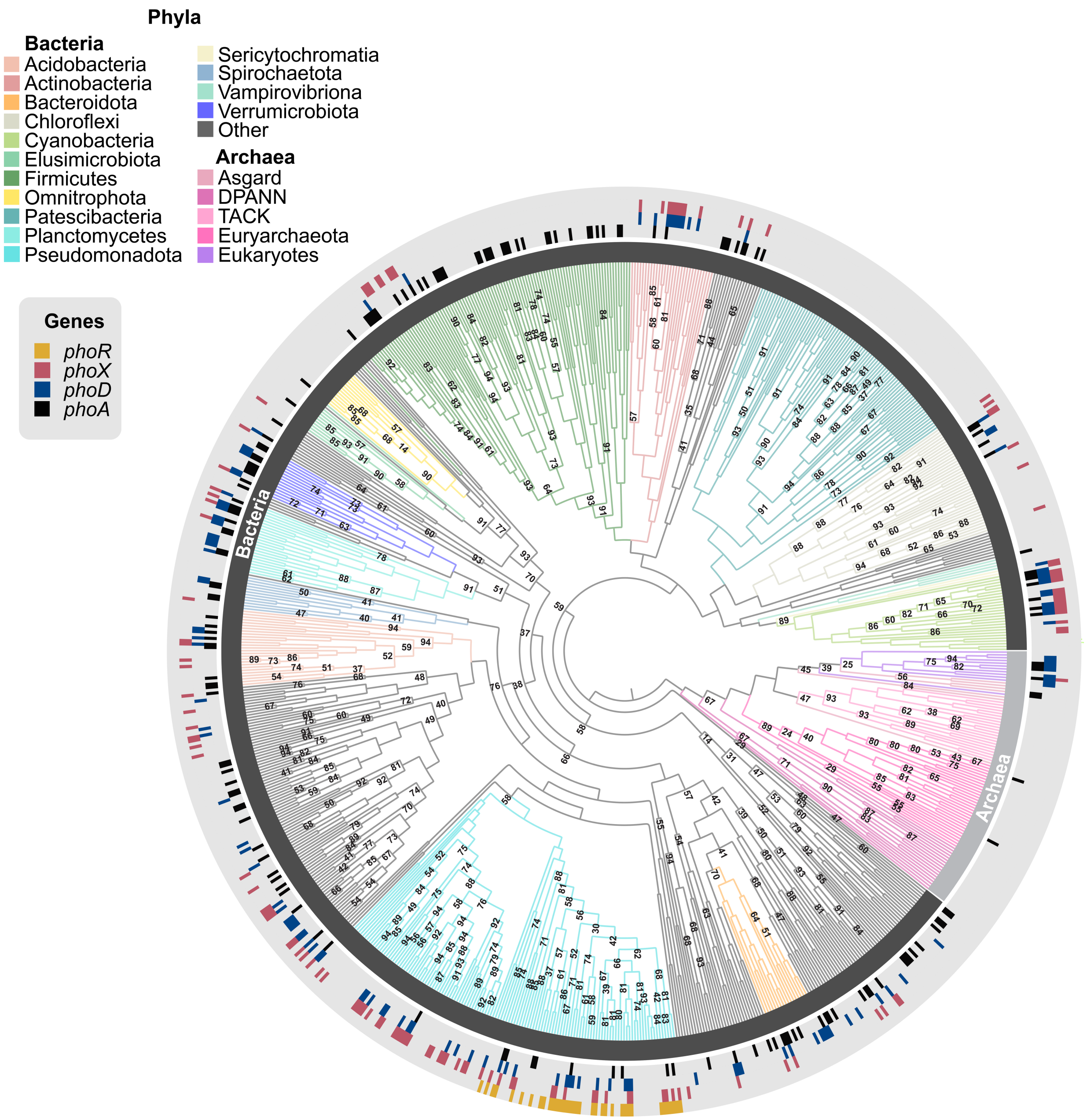


**Figure S2.** Distribution of alkaline phosphatase related genes in the tree of life, with associated support values. The presence of the genes *phoA* (black), *phoD* (blue), *phoX* (red), and *phoR* (yellow) are represented at the tips of branches (e-value threshold 0.001). The branches are colored according to their phylum. The phylogenetic tree was reconstructed from 16 ribosomal proteins using a maximum likelihood approach. Ultrafast bootstrap support values are labelled at branch points.

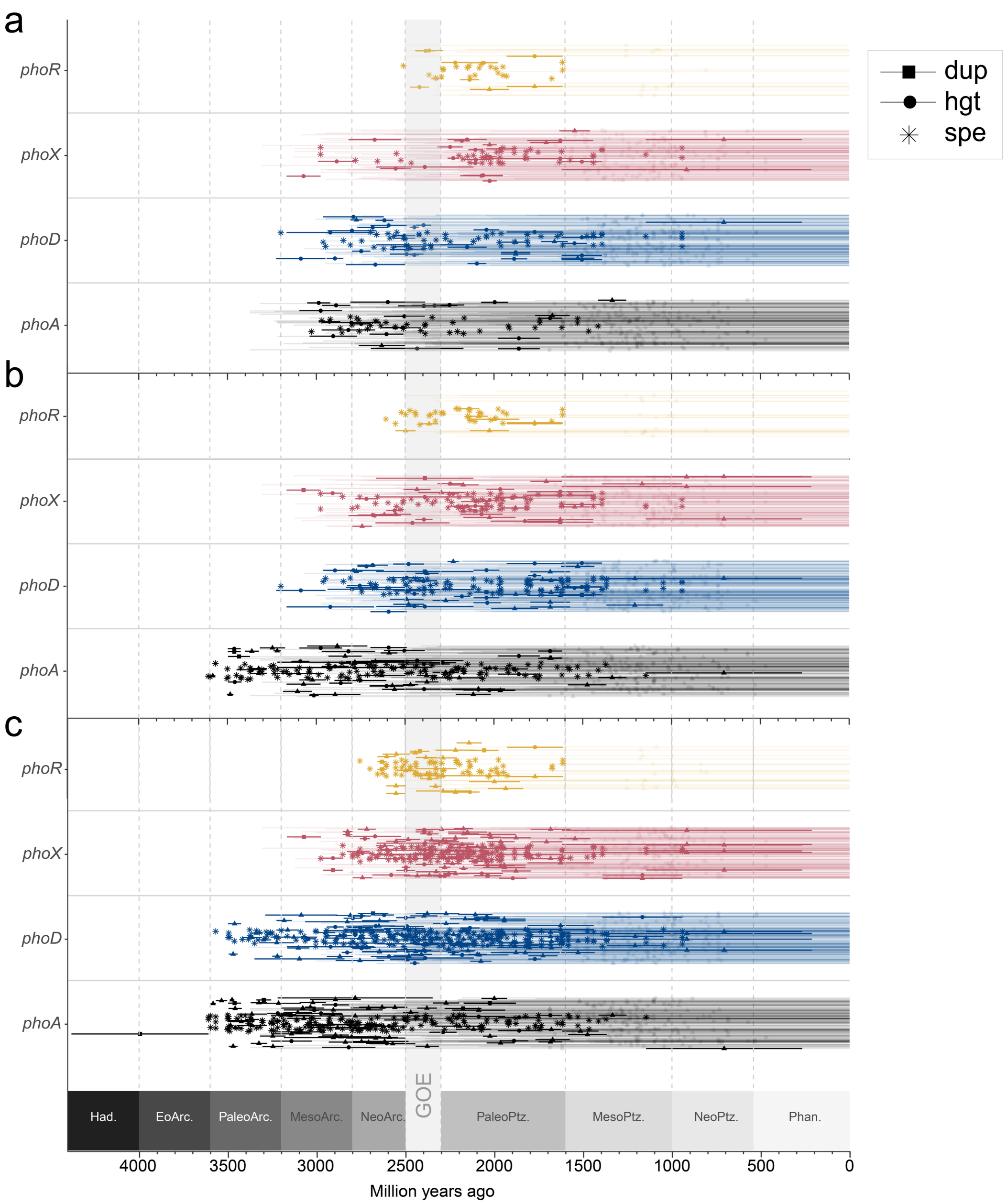


**Figure S3**. Effect of different costs for horizontal gene transfers on the estimated timing of evolution of *phoA*, *phoD*, *phoX*, and *phoR* using the CIR (Cox-Ingersoll-Ross) clock model. Reconciliations were performed with horizontal gene transfer costs of 2 (a), 4 (b), and 6 (c) using results of the molecular clocks made with the CIR clock model. Horizontal gene transfers (circles) and duplications (squares) of each gene are marked as the midpoints of lineages in which they are estimated to have occurred. Speciation events, marking a point in time where the gene was inherited from its parents, are marked with asterisks. Transparency indicates whether an event occurred on an internal (opaque) or terminal (faded) branch. GOE refers to the Great Oxidation Event.

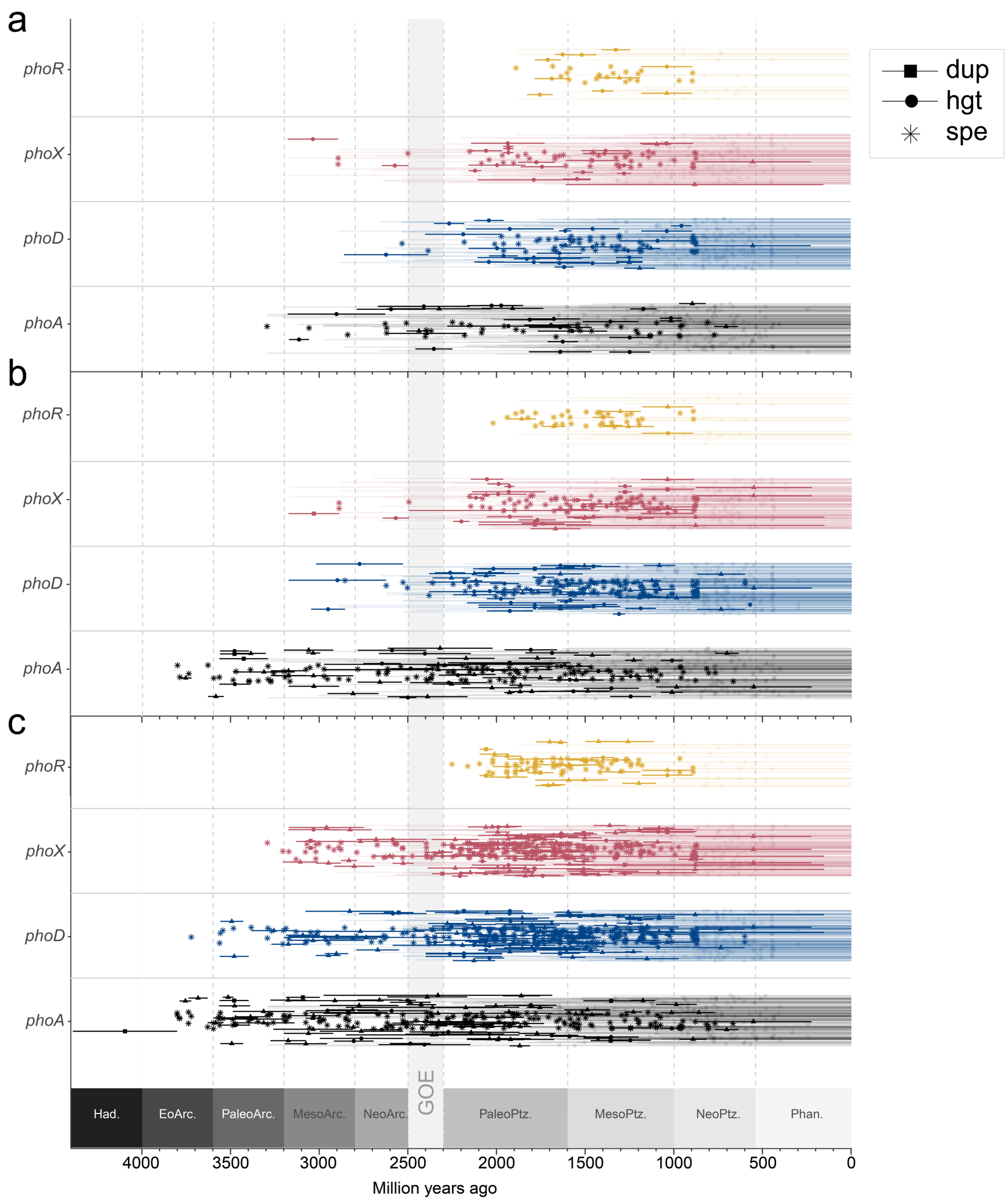


**Figure S4**. Effect of different costs for horizontal gene transfers on the estimated timing of evolution of *phoA*, *phoD*, *phoX*, and *phoR* using the LN (log-normal) clock model. Reconciliations were performed with horizontal gene transfer costs of 2 (a), 4 (b), and 6 (c) using results of the molecular clocks made with the LN clock model. Horizontal gene transfers (circles) and duplications (squares) of each gene are marked as the midpoints of lineages in which they are estimated to have occurred. Speciation events, marking a point in time where the gene was inherited from its parents, are marked with asterisks. Transparency indicates whether an event occurred on an internal (opaque) or terminal (faded) branch. GOE refers to the Great Oxidation Event.

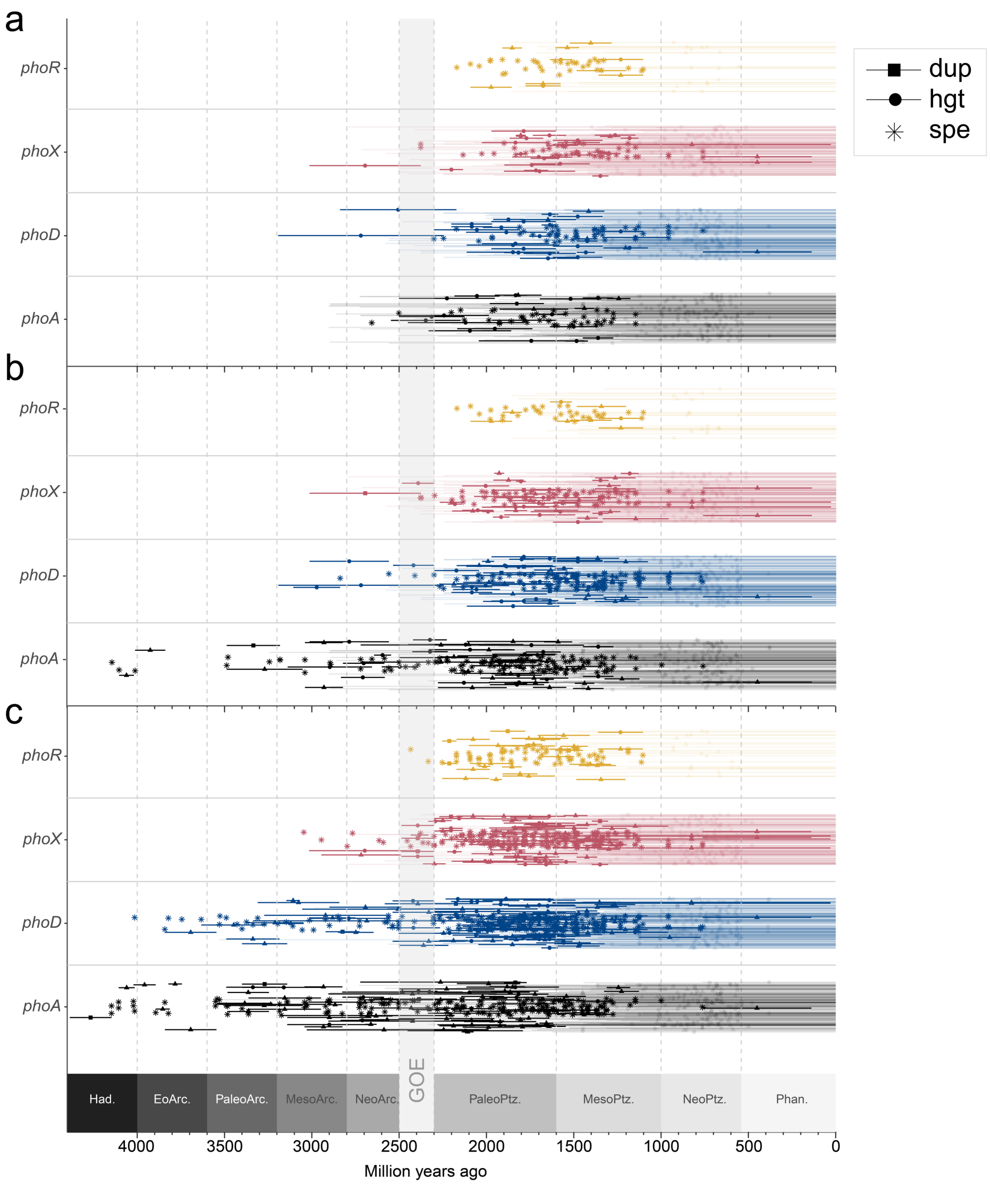


**Figure S5**. Effect of different costs for horizontal gene transfers on the estimated timing of evolution of *phoA*, *phoD*, *phoX*, and *phoR* using the UGAM (uncorrelated gamma multiplier) clock model. Reconciliations were performed with horizontal gene transfer costs of 2 (a), 4 (b), and 6 (c) using results of the molecular clocks made with the UGAM clock model. Horizontal gene transfers (circles) and duplications (squares) of each gene are marked as the midpoints of lineages in which they are estimated to have occurred. Speciation events, marking a point in time where the gene was inherited from its parents, are marked with asterisks. Transparency indicates whether an event occurred on an internal (opaque) or terminal (faded) branch. GOE refers to the Great Oxidation Event.

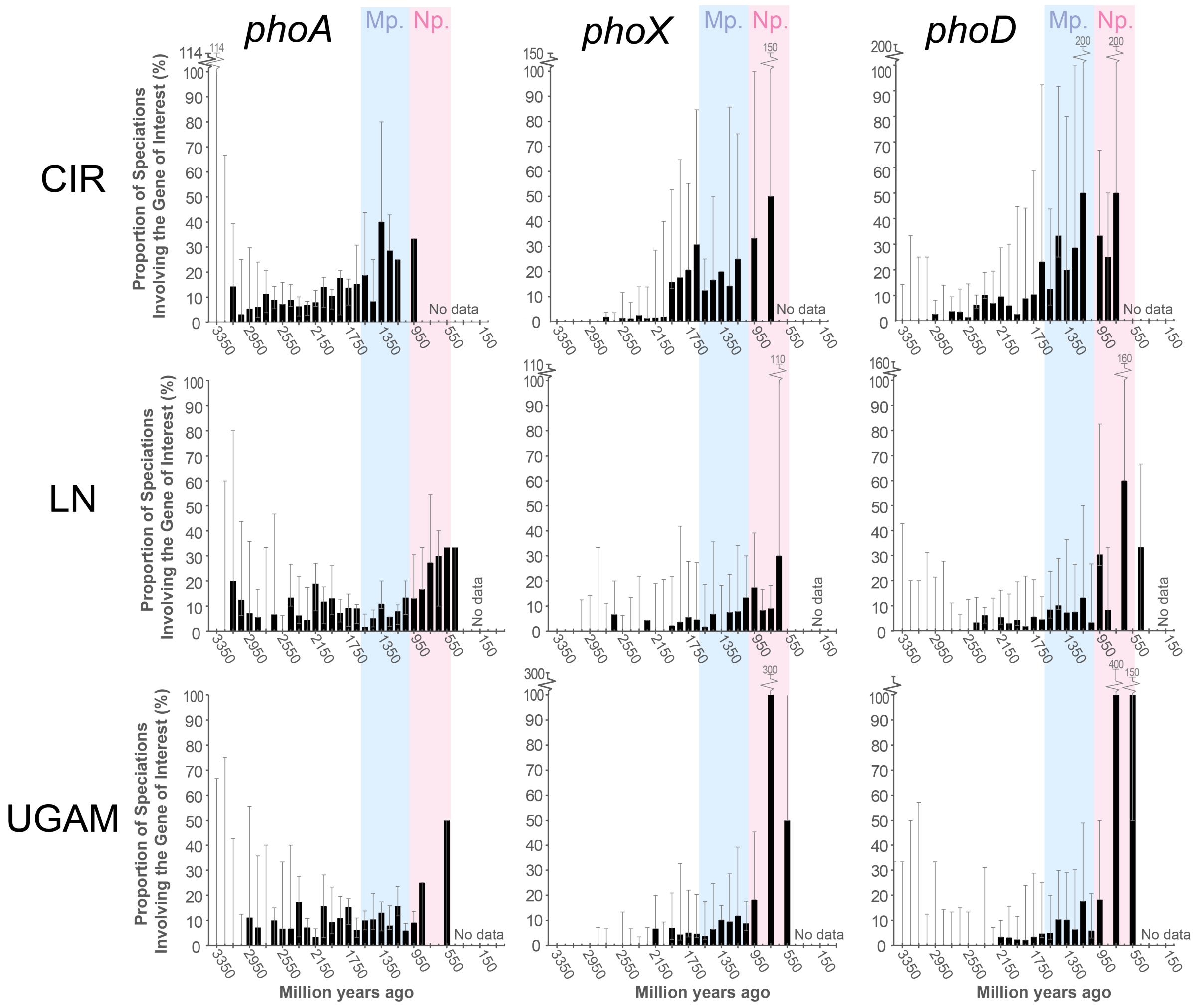


**Figure S6.** Proportional spread of alkaline phosphatase genes through time. Black bars represent the proportion of all speciation events which involved alkaline phosphatase genes calculated based on estimations made with the default horizontal gene transfer (HGT) cost (of 3). Error bars represent the range of values estimated using HGT costs of 2, 4 and 6. Blue shading highlights the Mesoproterozoic (Mp.) and pink shading highlights the Neoproterozoic (Np.). Values on the x axis indicate the midpoint of time bins spanning 100 million years. Graphs commence at 3550 million years ago, the mid-point of the oldest time bin containing at least 3 total speciations in all clock models (CIR, LN and UGAM).

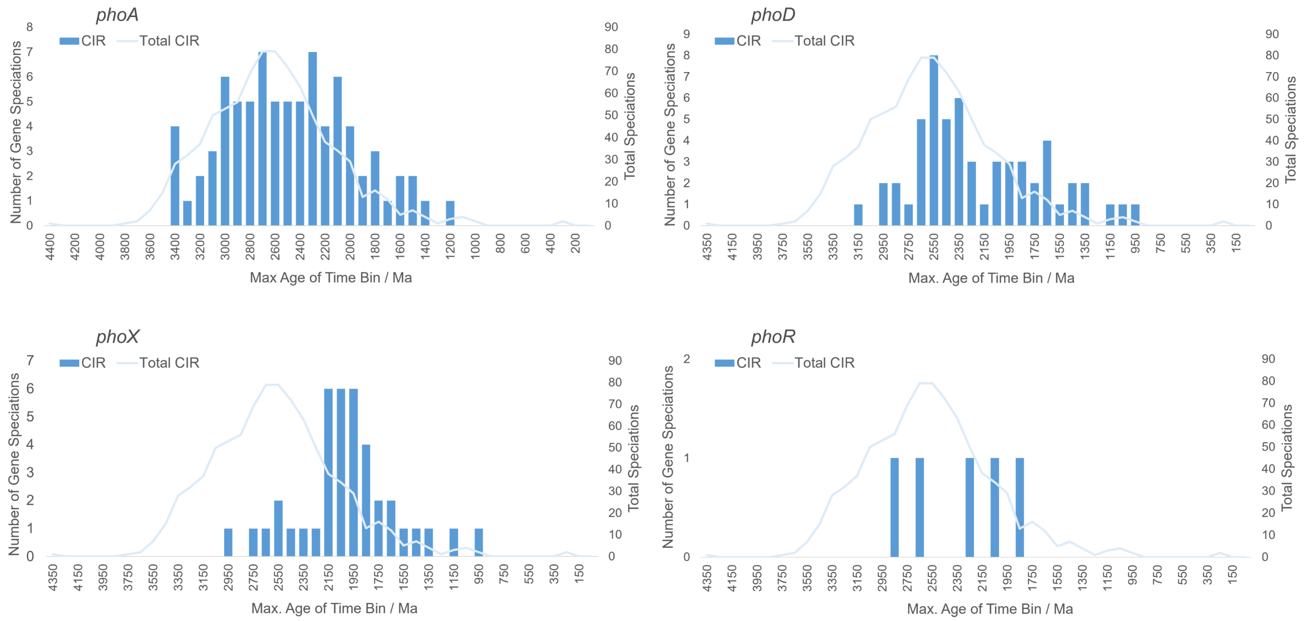


**Figure S7.** Number of speciation events in the time-calibrated species tree (line plotted on right y axis) and the number of speciation events involving *phoA*, *phoD*, *phoX*, and *phoR* (bars plotted on left y axis) through time. Values on the x axis represent the middle of 100-million-year time bins. Estimates made using the CIR clock model and a horizontal gene transfer cost of 3.

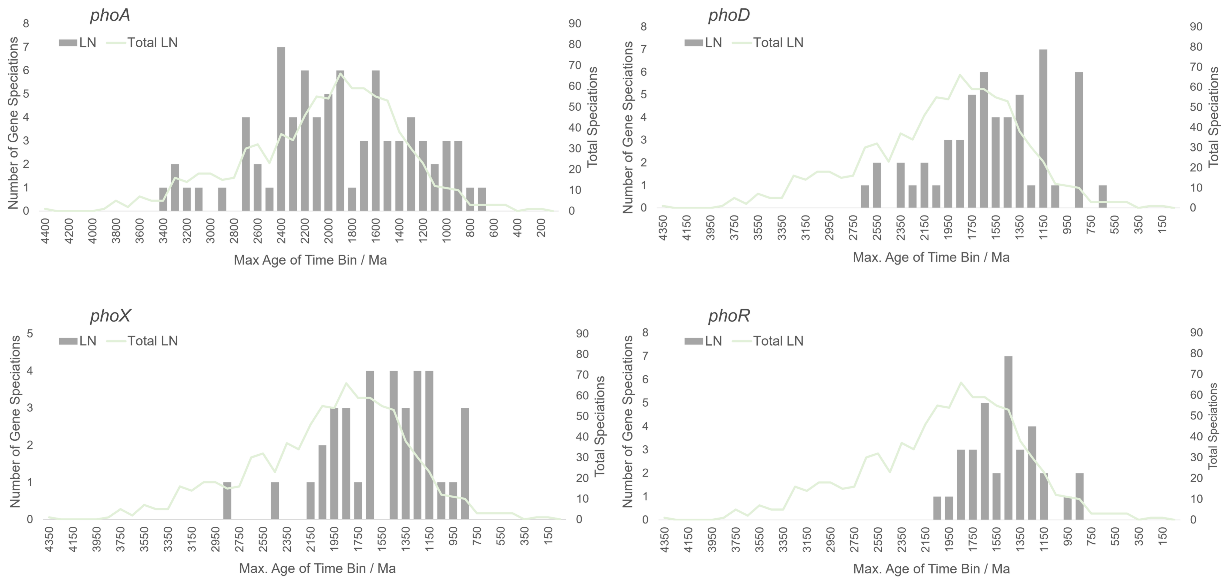


**Figure S8.** Number of speciation events in the time-calibrated species tree (line plotted on right y axis) and the number of speciation events involving *phoA*, *phoD*, *phoX*, and *phoR* (bars plotted on left y axis) through time. Values on the x axis represent the middle of 100-million-year time bins. Estimates made using the LN clock model and a horizontal gene transfer cost of 3.

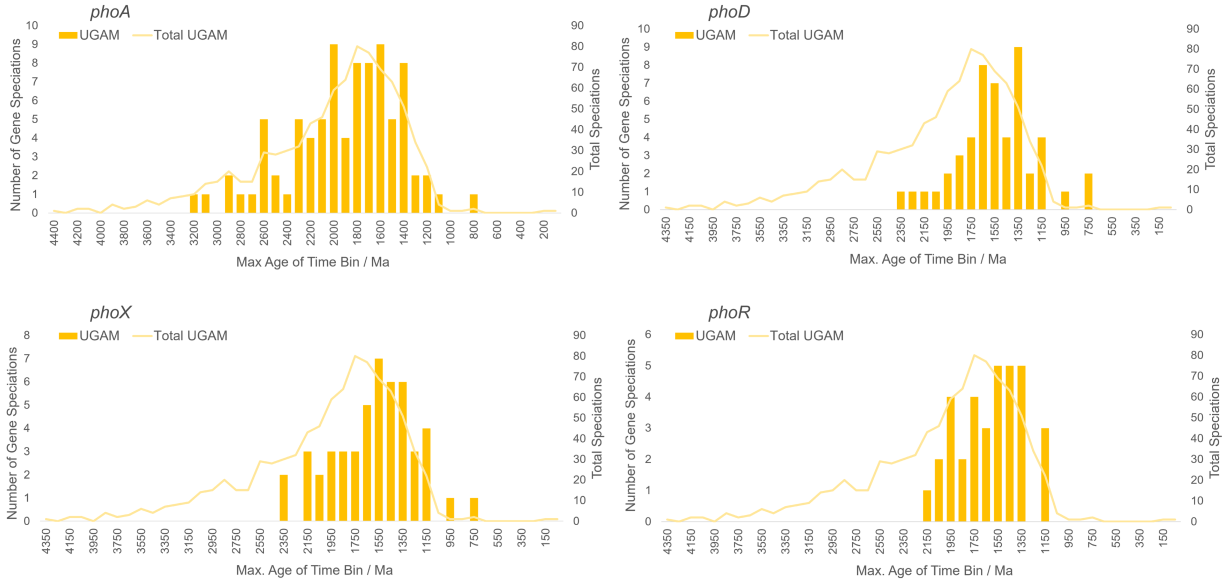


**Figure S9.** Number of speciation events in the time-calibrated species tree (line plotted on right y axis) and the number of speciation events involving *phoA*, *phoD*, *phoX*, and *phoR* (bars plotted on left y axis) through time. Values on the x axis represent the middle of 100-million-year time bins. Estimates made using the UGAM clock model and a horizontal gene transfer cost of 3.
